# Intraspecific encounters can induce home-range shifts

**DOI:** 10.1101/2023.06.07.544097

**Authors:** William F. Fagan, Ananke Krishnan, Qianru Liao, Christen H. Fleming, Daisy Liao, Clayton Lamb, Brent Patterson, Tyler Wheeldon, Ricardo Martinez-Garcia, Jorge F. S. Menezes, Michael J. Noonan, Eliezer Gurarie, Justin M. Calabrese

## Abstract

Direct encounters, in which two or more individuals are physically close to one another, are a topic of increasing interest as more and better movement data become available. Recent progress, including the development of statistical tools for estimating robust measures of changes in animals’ space use over time, facilitates opportunities to link direct encounters between individuals with the long-term consequences of those encounters. Working with movement data for coyotes (*Canis latrans*) and grizzly bears (*Ursus arctos horribilis*), we investigate whether close intraspecific encounters were associated with spatial shifts in the animals’ range distributions, as might be expected if one or both of the individuals involved in an encounter were seeking to reduce or avoid conflict over space. We analyze the movement data of a pair of coyotes in detail, identifying how a shift in home range location resulting from altered movement behavior was apparently a consequence of a close intraspecific encounter. With grizzly bear movement data, we approach the problem from the perspective of a set of encounter pairs within a population. We find support for the hypotheses that 1) close intraspecific encounters between bears are, on average, associated with subsequent shifts in range distributions and 2) encounters defined at finer spatial scales are followed by greater changes in space use. Our results suggest that animals can undertake long-term, large-scale spatial shifts in response to close intraspecific encounters that have the potential for conflict. These results lend support for existing theory on the evolution of territories and space use (e.g., Maynard-Smith’s bourgeois strategy regarding low-conflict coexistence). Overall, we find that analyses of movement data in a pairwise context can 1) identify distances at which individuals’ proximity to one another may alter behavior and 2) facilitate testing of population-level hypotheses concerning the potential for direct encounters to alter individuals’ space use.

**Open Research Statement:** Movement data for the coyotes and grizzly bears are posted on Movebank.org as datasets 1614661371 and 1044288582, respectively. Statistical tools for estimating, manipulating, and comparing home ranges from movement data are implemented in the open-source R package *ctmm*. R scripts used to carry out specific analyses for this study are openly available on GitHub at https://github.com/anagkrish/encounter_homerangeshift.

## Introduction

The boundaries of an animal’s home range can be affected by many factors (Tucker et al. 2014), including physical features (waterbodies, highways), food resources (Corriale et al. 2013, Pletenev et al. 2021), social systems (Harestad and Bunnel 1979), and escape cover and travel routes (Powell 2000). Among these, perhaps the most important factor determining territoriality is the presence of nearby conspecifics (Brashares et al. 2010, Schoepf et al. 2015). Species might actively defend their territories (Powell 2000), avoid areas in which they have a high probability of encountering neighboring individuals (Mech and Harper 2002), or be more alert when they move through possible encounter areas (Tórrez-Herrera et al. 2020). The importance of intraspecific interactions is well demonstrated by mechanistic home range analysis, which accurately predict spatial conformation of individual home ranges by modelling the impact of indirect interactions, such as scent marking, on the formation, structure and maintenance of home range boundaries (Moorcroft et al. 2006, Moorcroft and Lewis 2013). That these deterministic, interaction-based models can accurately predict the spatial conformation of individual home ranges highlights the importance of interactions in governing patterns of space use (Ellison et al. 2020).

Direct encounters are relatively rare and difficult to study, but are a topic of increasing interest in spatial ecology as more and better movement data become available (Courbin et al. 2016, Jordan et al. 2017, Broekhuis et al. 2019, Cheraghi et al. 2019, Périquet et al. 2021). Direct encounters between individuals also play a central role in evolutionary theory regarding territoriality, including considerations of the influence of cost-avoidance on behavior (Maynard-Smith 1982, Sherratt and Mesterton-Gibbons 2015). Tests of such ideas have historically been difficult to evaluate on the spatial scales typical of the home ranges of large mammals. This difficulty stems from the joint needs to first, identify that an encounter between individuals has taken place, and second, observe individuals long enough both before and after the encounter to have a clear view of the consequences. Relatively recent developments in movement ecology, such as the widespread availability of movement tracks with high temporal resolution (Nathan et al. 2022) and the development of statistical tools for estimating robust measures of changes in space use over time (Silva et al. 2022), hold promise for resolving this knowledge gap. Ecologists are now in a better position to identify encounter events, quantify the impacts of encounters on the individuals involved, and interpret those results in the context of theoretical predictions regarding space use.

Some work in this direction has already begun. For example, working in a landscape with several species of mammalian carnivores, Ruprecht et al. (2022) demonstrated how scavenging at kill sites increased opportunities for interspecific encounters, resource-transfer, and mortality events. Klauder et al. (2021) and Périquet et al. (2021) also explored encounters between individual predators in association with carrion, with the latter study discussing how encounters between individuals of different carnivore species at carrion or waterholes may lead to local-scale displacement of one or more of the individuals involved in the encounter. Noonan et al. (2021) developed methods for analyzing animal movement data to identify the locations within a set of nearby home ranges at which individuals were likely to encounter one another and further demonstrated that known encounters fell within zones of heightened encounter probability.

Several models have examined the likelihood and location of encounters taking place as a function of animal movement behaviors. For example, Gurarie and Ovaskainen (2012) examined scenarios involving encounters between foragers and stationary prey, leading to a taxonomy of encounter processes. Laidre et al. (2012) identified sex-based differences in the movement patterns of polar bears, finding that males’ more tortuous movements reduced the rate of male– male encounters, while having little impact on male–female encounters. Martinez-Garcia et al. (2020) analyzed stochastic models of pairs of moving animals, demonstrating how range-residency and a non-local perceptual range can alter the probability of pairwise encounters.

Despite progress on both theoretical and empirical fronts, much remains to be explored concerning the relationship between direct encounters and home range usage in animals (Noonan et al. 2021). Here, we examine the connection between close intraspecific encounters and home range dynamics. Working with data for two species of mammalian carnivores, we investigate whether close intraspecific encounters were associated with any spatial shifts in range distributions. We analyze the movement data of one such encounter pair in detail, identifying how a shift in home range location resulting from altered movement behavior was apparently a consequence of a close intraspecific encounter. In a separate analysis, we approach the problem from the perspective of a set of encounter pairs within a population, allowing us to 1) test the hypothesis that close intraspecific encounters are on average associated with shifts in range distributions and 2) understand how sex and seasonality influence the spatial consequences of such encounters.

These efforts apply a series of innovative statistical tools to detailed animal movement data, providing a rare opportunity to evaluate empirical evidence for longstanding—but little explored—hypotheses concerning mechanisms for territorial coexistence summarized in the ‘bourgeois’ and ‘anti-bourgeois’ strategies of Maynard-Smith (1982). These strategies highlight the relative utility of conflict avoidance and aggression, respectively, as routes to securing sufficient resources while minimizing risk (Maynard-Smith 1982, Morrell and Kokko 2005, Sherratt and Mesterton-Gibbons 2015, Hare et al. 2016).

## Materials and Methods

### Tracking data

To investigate evidence for shifts in animal home ranges before and after direct encounters, we used available GPS tracking data from Movebank.org (Wikelski and Kays 2022). We considered two datasets, each for a different purpose. First, for a detailed analysis of movement and space use associated with an encounter in a dataset with high temporal resolution for an extended duration, we considered movement tracks for a pair of coyotes (*Canis latrans)* from Ontario, Canada (Wheeldon 2020) (Table S1). Second, to illustrate how movement data for a large number of individuals in a population can be used to investigate whether encounters were, on average, associated with altered patterns of space use, we used a hypothesis testing framework at the set-of-pairs level, and considered data for N = 32 grizzly bears (*Ursus arctos horribilis*) living near Fernie in southwestern Canada near the border of British Columbia and Alberta (Lamb et al. in review) (Table S2). Grizzly bears are generally considered non-territorial in the sense that they do not actively defend spatial boundaries between individuals and allow spatial overlap with conspecifics at times (McLoughlin et al. 2000). In what follows, we explore the connection between encounters and space use, without any assumptions about territorial defense.

### Identification of Encounters and Estimation of Range Distributions

Working with pairs of tracks, we used the *distances*() function in the R package *ctmm* (Calabrese et al. 2016) to estimate distances between pairs of individuals that resided near each other in the same geographic area over an extended period of time. The function *ctmm::distances*() is a robust distance estimation method that accounts for, and is insensitive to, mismatched and irregular sampling between tracks as well as location error. For our core analyses, two individuals were said to “encounter” one another if they came within 100m of each other, and the time of the encounter was defined as the time at which there was the shortest distance between the pair of animals involved in the encounter. Our 100m threshold for defining encounters is arbitrary, but it is highly conservative compared to previous studies of carnivore encounter dynamics (800m in Jordan et al. 2017; 500m in Broekhuis et al. 2019 and Courbin et al. 2016; and 200m in Périquet et al. 2021). In a comparative analysis, we also considered encounter thresholds ranging from 50m to 500m.

Once encounters were identified, we fit a series of continuous-time movement models to the tracking data for each individual in each pair before, and separately, after, the encounter. The fitted models included the Independent and Identically Distributed (IID) process, which features uncorrelated positions and velocities; the Ornstein-Uhlenbeck (OU) process, which features correlated positions but uncorrelated velocities (Uhlenbeck and Ornstein 1930); and an OU-Foraging (OUF) process, which features both correlated positions and correlated velocities (Fleming et al. 2014a,b). We used AIC_c_-based model selection to identify the best model. Individuals that had an IID selected model were rechecked and refit; three individuals were dropped because they had very sparse sampling (fewer than 20 relocations).

We then used autocorrelated kernel density estimation (AKDE; Fleming et al. 2015, 2017, Silva et al. 2022) to estimate each individual’s home range before versus after the encounter as the corresponding range distribution (RD) and uncertainty of that distribution for the best-fit movement model. As modern high-resolution animal tracking data have become increasingly available, traditional methods of home range estimation, such as calculation of minimum convex polygons or conventional kernel density estimation, have been demonstrated to be unsuitable (Noonan et al. 2019). AKDE is a statistically efficient method for estimation of home ranges of animals whose movement data includes autocorrelation, small sample size, and missing or irregular data (Silva et al. 2022).

Having obtained four RD estimates for each encounter (i.e., before and after RDs for each individual in each pair), we then calculated differences between RDs using both the Bhattacharyya Distance (Bhattacharyya 1943, Winner et al. 2018) and the proportional overlap. The Bhattacharyya Distance (BD) is a unitless measure of dissimilarity between two probability distributions that takes values of zero for identical distributions and infinity for distributions with no shared support, and is readily incorporated into statistical tests. We also report results in terms of the Bhattacharyya Coefficient (BC) or ‘proportional overlap’, which is zero for distributions with no shared support and 1 for identical distributions. Overall, BD is more amenable to statistical analyses, but BC (proportional overlap) is more easily interpretable on both conceptual and visual grounds. Comparisons of individual distances, estimation of home ranges, determination of BDs, and calculation of RD proportional overlaps were conducted using *ctmm*. New code in *ctmm* created for this project allows for the propagation of uncertainties in calculating population-mean BDs; these updates appear in *ctmm* version 1.1.0 involving the functions *overlap()* and *meta()*.

### Pair-level analysis: Coyotes

We identified a pair of coyotes that demonstrated a close (66 ± 1 m) encounter, which was about 1 order of magnitude closer than the animals came to each other during the 8 months that they were simultaneously tracked. We then investigated aspects of the movement tracks and the spatial consequences that could be attributed to this encounter in detail.

We estimated home ranges for each individual before and after the encounter, and determined the degree of spatial overlap between RDs as described above. To provide further support for a shift in home ranges as a consequence of the encounter, we estimated home range shifts at other times when the animals were actually far apart, which we termed “null encounters.” These null encounters were selected to span even intervals across the period over which both animals were tracked. Home ranges for each individual and RD overlaps were calculated before and after each null encounter as described above. This allowed us to verify that any change in RD overlap was occurring uniquely following the encounter and not independent of encounter distance or at other times as well.

Lastly, we calculated the ballistic length scale for each individual as a running measure across 60 days of movement data. Ballistic length scale (denoted *l*_*ν*_) is a measure of linearity in movement (Visser and Kiørboe 2006, Noonan et al. 2023) and is calculated by:

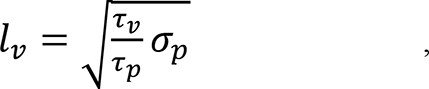

where the parameters *τ*_*ν*_ (timescale of autocorrelation in velocity; seconds), *τ*_*p*_ (timescale of autocorrelation in position; seconds), and *σ*_*p*_ (spatial variance in movement; m^2^) are estimated via the *ctmm* package from an OUF model for range-resident movement (which was the best-fit model for these individuals). This allowed us to quantify the directedness of each individual’s movement before and after the encounter.

### Population-level analysis: Grizzly Bears

We next tested the hypothesis that spatial shifts following encounters are consistent features of a population of grizzly bears. Range distributions (RDs) and spatial shifts associated with encounters were calculated for each pair of bears that came within 100m of each other (N = 44 pairs, which includes two bear-pairs with encounters in different years). We tested the statistical significance between RD overlaps before and after encounters on the whole population using a novel *χ^2^* inverse-Gaussian (*χ^2^*-*IG*) meta-analysis framework (Fleming et al. 2022). This framework employs a non-linear hierarchical model developed for estimating population-mean home-range parameters from individual home-range estimates, while propagating their uncertainties (Fleming et al. 2022). The methods, now incorporated into the R package *ctmm*, model population-level estimates as having an inverse-Gaussian distribution, which is appropriate for individual scale parameter estimates, like home-range size and distance, with large corresponding uncertainties. Analyses return a statistical comparison between models assuming that the before-encounter and after-encounter RDs are (or are not) distinguishable. The difference between RDs is measured via the BD, and model support is scored via AIC_c_. The grizzly bear data were not amenable to analyses of null encounters or ballistic length scale calculations because the grizzly bear tracks were relatively shorter in duration and because the encounters often occurred within only a few weeks of the end or onset of hibernation.

## Results

Basic data on the movement tracks for coyotes PEC068 and PEC088 appear in Table S1. The coyotes encountered each other around 3:01 AM on 26 May 2012, coming within 66m of each other. This is by far the closest they came to each other during an ∼8-month window of simultaneous tracking (median distance: 7249 m, IQR: 5225-9056 m; Fig. 1A).

**Figure 1.**
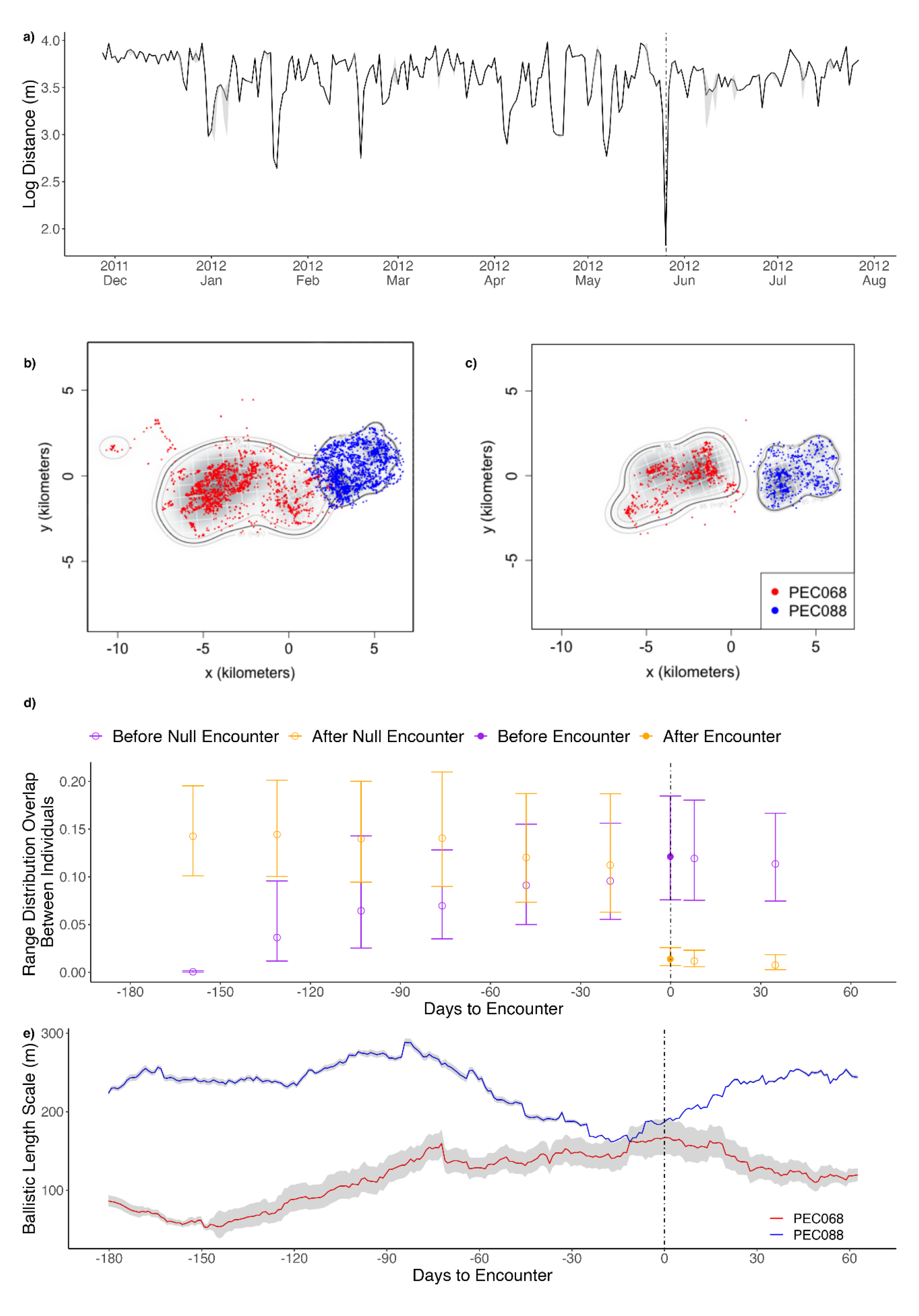
Detailed analyses of a close encounter between two coyotes. Inter-individual distances between coyote individuals PEC068 and PEC088 in Ontario, Canada, over an ∼8-month period in 2011–2012 (Panel A). When missing data necessitated interpolation of coyote positions, estimated inter-individual distances (+/- 95% CI as gray shading) are plotted. GPS tracking data indicate that the coyotes’ movements brought them within 66m of each other on 26 May 2012 (shown as day 0). Panels B and C plot home ranges (+/- 95% CI) for the coyotes before and after this encounter, respectively, calculated as range distributions in the R package *ctmm*. Note the spatial shift by PEC068, reducing overlap with PEC088 after versus before the encounter (ΔAICc = 24.04). Panel D provides the proportional overlap (Bhattacharyya Coefficient +/- 95% CI) of the individuals’ home ranges before (purple) and after (orange) the actual encounter (denoted by solid symbols and the dashed vertical line) compared with similar overlaps measured, for comparative purposes, for alternative ‘null-encounter’ dates when encounters did not occur (open symbols). Panel E gives each individual’s ballistic length scale (+/- 95% CI) calculated on a running basis for 60-day windows. Note that the ballistic length scale of PEC068 (whose home range shifted following the encounter, thus decreasing inter-individual home range overlap) decreased by ∼50% in the 60 days after the encounter whereas that of PEC088 increased by ∼60%.

In the months prior to this encounter, the coyotes’ home ranges overlapped by 12.1% (Fig. 1B); following the encounter, PEC068 shifted and shrank its home range by ∼30%, reducing the overlap to only 1.1% (Fig. 1C). This reduction in home range overlap is significant: a model in which the RDs have shifted so as to show less overlap after versus before the encounter is better supported by 24.04 ΔAICc units versus a model in which the before versus after distributions are not distinguishable. In contrast, a selection of alternative ‘null encounter’ dates preceding the observed encounter yielded no evidence of home range shifts (Fig. 1D). In the 60 days after the encounter, the ballistic length scale (which is calculated over a 60d running window, see Methods) for PEC068 decreased by ∼50% whereas that for PEC088 increased by ∼60%.

Pairwise analyses of encounters between grizzly bears also demonstrated shifts in home ranges following encounters. When an encounter between a pair of bears occurred within <100m, the encounter was, on average, associated with changes in movement that increased the BD between the bears’ home ranges following the encounter (Fig. 2). Encounter-related shifts in home ranges were significant for the entire set of 44 bear pairs as a whole, but encounters between bears during the late fall contributed most strongly to this effect.

**Figure 2.**
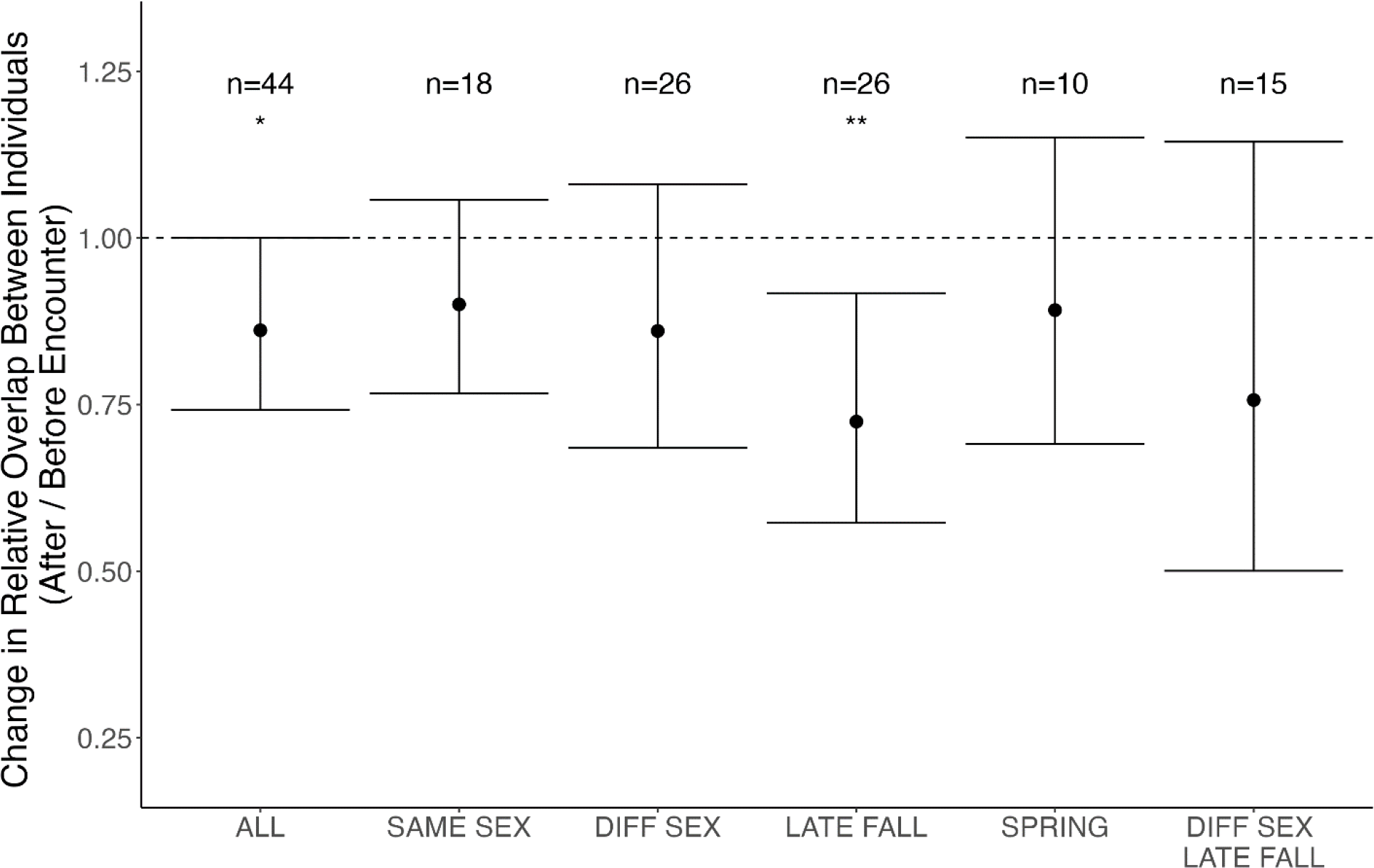
Mean (+/- 95% CI) relative change in home range overlap between pairs of Canadian grizzly bears that encountered each other at distances <100m. The y-axis plots the ratio of the home range overlap after an encounter versus home range overlap before the encounter; thus, values less than 1 indicate decreases in pairwise overlap. Results are shown for all pairs of bears exhibiting an encounter, all same-sex pairs, all different-sex pairs, all late fall encounters (1 September – 30 November), all spring encounters (1 June – 30 July), and all different-sex pairs whose encounters occurred during late fall. Asterisks above the error bars indicate significant changes in overlap: * p < 0.05, ** p < 0.01.

In contrast to the strong and significant results presented above at the population-of-pairs level, clear demonstrations of before versus after shifts in home ranges at the level of individual pairs of bears proved more difficult. This was due to uncertainty inherent in the location of the estimated home range boundaries that was carried through to assessments of the BDs (and, similarly, the proportional overlap) between RDs. Figure S1 provides plots of proportional overlap of RDs between the individuals in each pair of bears before versus after an encounter. The male-female pairs of EVGM86-EVGF66 (2021), EVGM77-EVGF85 (2019), EVGM90-EVGF73 (2018), and EVGM86-EVGF109 (2020) showed the greatest decreases in proportional overlap of RDs from before to after the encounter, but even these would not be deemed statistically significant given the large uncertainties in proportional overlap (Fig. S1). Encounter-related home range shifts were also visualized in terms of the shifts in RDs within individuals in each pair before versus after an encounter to identify whether one bear shifted its range more than the other bear involved in a given encounter (Fig. S2).

Repeating the above analyses using distance thresholds of <50m, <200m, and <300m to define an encounter yielded similar results; however, with more liberal thresholds of 400m to 500m, we found no evidence for home range shifts following encounters (Fig. 3). For encounter thresholds of 50m to 300m, late fall encounters (and separately, late fall encounters between male and female bears), were associated with the largest changes in home range overlap. In some cases, cubs were known to be accompanying a female involved in an encounter, and in these cases, BDs between individuals’ RDs tended to be greater following the encounter than before, but the effect was not significant for any encounter thresholds.

**Figure 3.**
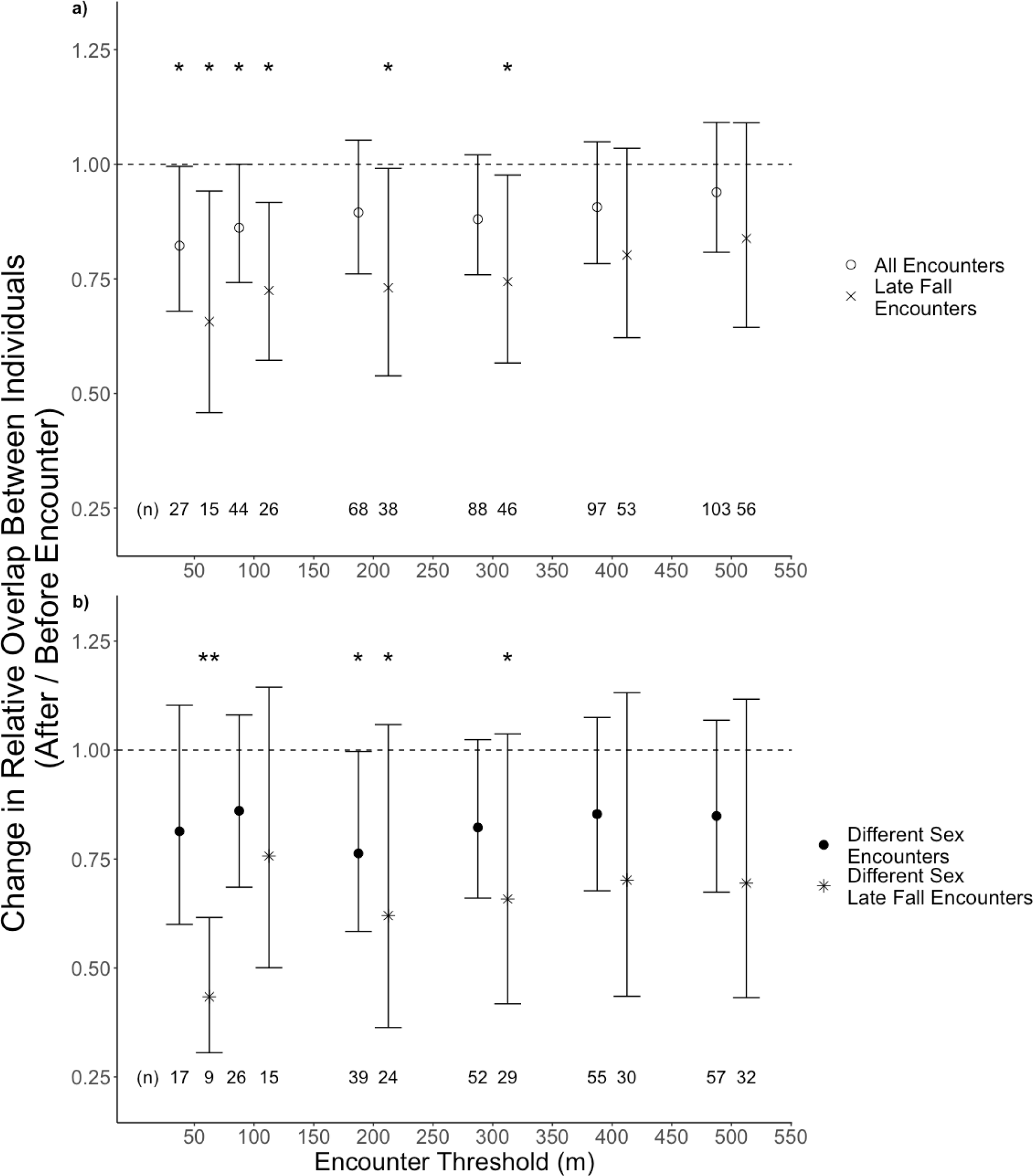
Mean (+/- 95% CI) relative change in home range overlap between pairs of Canadian grizzly bears for different definitions of what constitutes an encounter. The y-axis plots the ratio of the home range overlap after an encounter versus home range overlap before the encounter; thus, values less than 1 indicate decreases in pairwise overlap. Results in Panel A are shown for all pairs exhibiting an encounter and all pairs involving an encounter in late fall (1 September – 30 November). Panel B plots results for all different-sex pairs and all different-sex pairs involving encounters during late fall. Note that sample sizes in both panels depend on the definition of encounter distance. Asterisks above the error bars indicate significant changes in overlap: * p < 0.05, ** p < 0.01.

Encounters between pairs of bears, and especially encounters between male – female pairs, occurred far closer to carcass pits (locations where road kill and hunters’ gut dumps were systematically deposited) than did generic non-encounter fixes between pairs of bears (Fig. S3). Encounters between pairs of bears during the Late Fall period (1 September – 30 November) tended to occur closer to carcass pits than did generic non-encounter fixes during the same time period (Fig. S4).

## Discussion

Here we have demonstrated how encounters, defined on the basis of spatial proximity in pairwise analyses of GPS tracks for mammalian carnivores, can be associated with shifts in home ranges. Through a combination of detailed pairwise analysis and population-level hypothesis testing, we characterized changes in the overlap between individuals’ ranges and explored how sex and seasonality influence the spatial consequences of encounters.

### Coyotes

The coyote data suggest a scenario whereby the encounter yielded an outcome in which, for the remainder of the tracking period, one individual, PEC068, was a clear ‘loser.’ The range distribution of PEC068 decreased by ∼30% following the encounter, and that coyote no longer used the easternmost part of the range it occupied before the encounter (Fig 1). In the context of a larger, regional study of coyote home ranges and territoriality (Wheeldon 2020), PEC068 was determined to have exhibited an ambiguous space use pattern, and herein is considered a “non-territorial resident,” exhibiting relatively weak range fidelity and undertaking multiple forays outside of its range that intruded on territories of other coyotes, including but not limited to PEC088. As such, the before versus after encounter RDs estimated for PEC068 would correspond to ‘undefended home ranges’ rather than territories *per se*, and the spatial shift observed for this animal appears to have derived from a post-encounter reduction in its foray activity rather than contraction of a defended territory. The ballistic length scale results support the above interpretation in that PEC068 decreased the linearity of its movement following the encounter, presumably reflecting the reduction in its foray activity (i.e., out-and-back movements), whereas PEC088 increased the linearity of its movement following the encounter, which may partly reflect increased territorial behavior in the form of patrolling the perimeter (note the relative increase in the intensity of PEC088’s usage of the western part of its range following the encounter in Fig. 1c). The encounter occurred during the denning season and PEC088—based on its reproductive history—may have had pups at the time, which may have heightened the intensity of its encounter and subsequent activity.

### Grizzly Bears

When occurring at distances <300m, encounters between bears were, on average, associated with substantial shifts in RDs (Figs. 2, 3). However, when we broadened the definition of encounter to include proximity events occurring beyond this 300m encounter threshold, we found no evidence for a population-level association between encounters and RD shifts. Previous studies using animal movement data to investigate the consequences of encounters in other systems have relied upon a broad range of threshold distances, including some as great as 500m to 800m. Our results suggest caution in relying upon such large distances to delineate encounters between tracked individuals. Here, very loose definitions of encounters indicated the absence of encounter-associated changes in space use that were in fact observable with more restrictive assumptions (Figures 2, 3). Moreover, smaller distance thresholds when defining encounters were associated with stronger spatial changes, likely because those smaller thresholds were more apt to correspond to actual encounters (i.e., one or both individuals detecting the presence of the other). Experimental work involving human proximity to wolves presents similar ideas, and emphasizes that encounters can be decidedly one-sided in terms of detection and spatial response (Versluijs et al. 2022). We recommend that future encounter-based research conduct sensitivity analyses to identify the encounter distances that are most strongly associated with changes in space use. Such analyses could have the additional benefit of providing insight into just how big are the perceptual ranges of the animals involved (Zollner and Lima 1997, Boonman et al. 2013, Fagan et al. 2017).

Of all the categorizations we considered, encounters between male and female bears during the late fall resulted in the greatest shifts in home range overlap (Fig. 3). During this season, encounters between bears occurred disproportionately close to carcass pits (Fig. S4), and may involve heightened aggression during a period of increased resource competition in the weeks preceding hibernation.

### Encounters, range distribution shifts, and the evolutionary theory of territories

The occurrence of an encounter depends not just on proximity, but also on perceptual abilities (visual, olfactory, auditory) and spatial context. In the case of the coyotes, the close encounter occurred near the intersection between two fields separated by a hedgerow, so it is almost certain that at least one of the coyotes detected the other. Indeed, accelerometer data from around the time of the coyotes’ encounter (Wheeldon, unpublished data) suggest that the encounter may have been a ‘close-call’ for PEC068 rather than an encounter involving direct aggression. PEC068 was inactive for ∼50 minutes shortly after the encounter (∼ 10 minutes), indicating that it may have bedded down to hide from PEC088, which continued to be active for ∼35 minutes post-encounter and then became inactive, at which point PEC068 became active and eventually left the territory of PEC088. In the case of the bears, the encounters occurred primarily in broken timber such that proximity may not always have led to mutual detection, which could have contributed to the heterogeneity in pairwise results even for encounter thresholds of 100m (Figures 2, S1, S2).

Our results suggest that animals can undertake long-term, large-scale spatial shifts in response to close intraspecific encounters that have the potential for conflict. These results not only corroborate the use of home range overlap to quantify encounter potential, but also corroborate some of the existing theory on the evolution of territories and space use. For example, in the case of the coyotes, our observation that only one individual continued to use the disputed area, over a period of more than three months following the encounter, indicates that “ownership” of this area is being respected by the losing party (PEC068). This form of low-conflict coexistence is known as the “bourgeois strategy” (Maynard-Smith 1982, Sherratt and Mesterton-Gibbons 2015). Theory has predicted that this strategy should be selected for in most natural environments, since it reduces the overall risk of injury for all parties involved (Maynard-Smith 1982, Sherratt and Mesterton-Gibbons 2015). Further, theory also suggests that the “bourgeois strategy” should occur when animals have the liberty of reducing their activity while maintaining their energetic budget (Menezes and Oliveira-Santos 2021). Considering that coyotes can substantially vary their daily activity and diet (Kitchen et al. 2000), PEC068’s reduction in foray activity that resulted in decreased overlap with the territory of PEC088 (Fig. 1), need not necessarily have translated into a loss of resources. Combining this support with previous studies indicating that coyotes forage optimally (Hérnandez et al. 2002), there is mounting evidence suggesting current theory can contribute to the successful prediction of coyote behavior.

Evidence from the bears, on the other hand, provided support for theory in a different manner. The population-level increase in average dissimilarity of RDs following encounters suggest a reduction of contact that could lead to aggressiveness, but it is not clear if this reduction was mutual or one-sided. The bear data can only rule out one of the most aggressive strategies, known as “anti-bourgeois” or “paradoxical” strategy of coexistence (Maynard-Smith 1982, Morrell and Kokko 2005). In this strategy, individuals show no respect for ownership and become more aggressive if displaced. If that were the case, recurring contacts would yield heightened conflicts, but no change in overlap, conflicting with our findings (Figures 2, 3). Because heightened aggression and repeated conflicts pose dangers for animals’ well-being, many studies have proposed that the paradoxical strategy would quickly be dominated by a modified “bourgeois strategy” involving better communication between rivals or “draws” during disputes (Hare et al. 2016, Morrell and Kokko 2005, Sherratt and Mesterton-Gibbons 2015). Thus, both datasets provide support for existing models of territorial evolution, but with respect to different strategies.

## Conclusion

Ecologists broadly recognize the importance of encounters between individuals; however, encounters are hard to quantify from data, and their long-term consequences have proved difficult to study. Increasing availability of high resolution movement data, together with a host of analytical tools to estimate home ranges, develop robust measures of spatial overlap, calculate ballistic length scales, and evaluate spatial context, create vastly improved opportunities to both identify encounter events and quantify their impacts on the individuals involved. Encounter-based analyses can be used to interpret changes in space use, identify distances at which individuals’ proximity to one another may alter behavior, and test population-level hypotheses concerning the potential for direct encounters to alter individuals’ space use.

With approaches and computational resources like those illustrated here, future work could investigate to what extent multiple behavioral shifts are consistent within individuals (indicating winner/loser dynamics in the context of territorial interactions). For example, little-investigated theoretical issues, such as transitions between ideal free and ideal despotic distributions, could be investigated by connecting encounter data with assessments of resources lost and gained as a result of spatial shifts in occupied ranges.

## Acknowledgements

W.F.F., C.H.F., and J.M.C. were supported by NSF IIBR 1915347. The University of Maryland provided additional financial support. R.M-G. was supported by FAPESP BIOTA Young Investigator Research Grant no. 2019/05523-8457 and the Simons Foundation through grant no. 284558FY19. This work was partially funded by the Center of Advanced Systems Understanding (CASUS), which is financed by Germany’s Federal Ministry of Education and Research (BMBF) and by the Saxon Ministry for Science, Culture, and Tourism (SMWK) with tax funds on the basis of the budget approved by the Saxon State Parliament.

## Author Contributions

W.F.F., Q.L., and C.H.F conceptualized the study. A.K., Q.L., and D.L. conducted statistical analyses and visualization under guidance from W.F.F, with J.M.C. and C.H.F. providing additional suggestions regarding the statistical analyses. B.P., T.W, and C.L. provided data and biological insights. C.H.F. developed new computer code now included in the open-source R package *ctmm*. W.F.F. wrote the initial draft with input from A.K., Q.L., and J.S.F.M. All authors contributed to interpretation of results plus review and editing of later stages of the draft.

